# Assessing fish welfare in small-scale commercial fixed-net fisheries off the southern Portuguese coast

**DOI:** 10.1101/2025.07.27.667075

**Authors:** Vighnesh Samel, Rita A Costa, Ana Marçalo, Magda Frade, Luís Bentes, João L. Saraiva, Jorge MS Gonçalves, Pedro M Guerreiro

## Abstract

Despite a growing interest in animal welfare in production systems, research on fish welfare remains limited, particularly in commercial fisheries. Fish caught in fixed-net fisheries experience multiple stressors from the time of capture to mortality on deck considered detrimental to their welfare. We examined the impact of bottom-set gill nets and on-board handling on catch welfare using behavioural and physiological indicators. Vitality assessments were performed on four commercially important fish species on-board fishing vessels through a devised vitality scale that included behaviours, morphological condition and reflexes as indicators of welfare. Physiological stress parameters (Cortisol, Glucose, Lactate and Osmolality) were evaluated in blood collected on deck and analysed in relation to the vitality scores. The vitality at arrival on deck as well as the rate of decrease in vitality differed significantly amongst the tested species. Furthermore, Generalised Linear Models predicted that several biological, operational, and environmental variables significantly affect the extent of time the fish shows activity, and hence, on the welfare. Elevated average cortisol levels were found at all the vitality stages highlighting the stress experienced by fish due to the fishing process. The findings of this study enable us to recommend welfare-friendly methods in set-net fisheries to promote better fishing standards.

## 1. Introduction

Welfare, broadly referring to the state of an individual in relation to its environment, is a topic of growing concern and importance. An individual is said to be in a state of ‘good welfare’ if it can cope with the environment, meeting the individual needs, while conversely, failure to cope or difficulty in fulfilling needs can be termed as ‘poor welfare’ [1,2]. Difficulties arise when applying such concepts to activities in which the ultimate goal is to harvest animals for human consumption, which invariably affect the welfare of the animals [3], and in this regard, the welfare of fish has generally been overlooked when compared to other vertebrates [4]. In recent years, attention has been given to farmed fish, particularly to the conditions of fish at harvest and slaughter [5–7], but little has been promoted to improve welfare during catch and subsequent processes in the fishing industry. This is an outstanding issue that includes ethical questions towards the animals, fair trade, food quality, and safety, and which deserves the consolidation of scientific evidence regarding future change in the industry standards.

Commercial fishing is a key source of food and income, and supports the livelihoods of a very large proportion of coastal communities, providing crucial nutrition and generating economic opportunities, particularly for small-scale fisheries [8]. However, it is important to recognise that fishing operations have a negative impact on fish welfare causing stress, physiological dysfunction and eventually injuries that accelerate mortality and reduce the survival of released or escaped individuals [9,10]. Thus, improving welfare is in fact based on changing methods to reduce or mitigate these negative impacts [9,11,12], with potential benefits on product quality. However, progress is limited by gaps in understanding how physiological stress arises throughout fishing operations and how it translates to welfare and survival post-release.

Historically, three approaches have been taken to study animal welfare. The nature-based approach entails that all animals have an inherent ‘nature’ and that animals in a good welfare state should be able to lead a natural life [13,14], while the feelings-based approach requires the animal in a good welfare state to ‘feel well’, free from the experiences of prolonged and intense pain, hunger, fear, and other negative states [15,16]. The latter, however, has been criticised in fish welfare studies since it assumes fishes have the capacity to have conscious subjective experiences, which are difficult to measure objectively [17,18]. Therefore, the present study was performed using the biologically defined function-based approach according to which good welfare implies that the animal is in good health and that its biological systems are performing well. This approach enables an objective way to measure welfare by measuring certain biological parameters [6,9]. Currently, no standardised protocol exists to measure fish welfare in capture fisheries. However, evaluating physiological parameters and its correlation with immediate or delayed mortality, can help identify welfare indicators [12,19]. Pre-mortality stress levels, particularly elevated cortisol, entail a low welfare state often accompanied by the imbalance of several other blood and cellular parameters [20–23]. Behavioral analyses, particularly vitality assessments, provide an operational approach to evaluate fish welfare, with lower vitality scores indicating higher stress levels [24–27]. Categorical Vitality Assessment (CVA) is a key method used to assess vitality, offering insights into pre-mortem stress experienced by fish during commercial bottom-set net fisheries [28].

Gill nets, whether set or drift, are one of the main métiers used by the small-scale artisanal fisheries, a sector responsible for about half of the wild-capture fisheries in the world production [29,30]. These nets catch the fish by their gills or entangle them in the netting, clearly impacting the welfare of target and non-target fish species [31,32], and continue fishing when lost, accounting for underestimated fish mortality [33]. In addition, these nets have a high incidence of immediate and delayed mortality in released/discarded fish [12]. Fish trapped in these nets may struggle for prolonged periods, experiencing various stressors throughout the fishing process, including injury, asphyxiation, hypoxia, exhaustion and osmoregulatory distress [4,9]. Several specific modifications in gear to reduce bycatch or facilitate disentanglement have been proposed [29,34]. However, it is important to note that reducing the stress during capture, while favouring the survival of released fish, may counterintuitively reduce the welfare of target species by increasing their survival time on-board, since presently the fish are not actively stunned and killed. Instead, they eventually die by asphyxiation on deck or during storage. Thus, these challenges identified during capture may persist during retrieval, on-board handling, and slaughter, posing significant welfare concerns.

The present study, conducted under a function-based approach of welfare [17,35,36], aimed at characterising the impact of bottom-set nets used in small-scale fisheries in the Southern Portuguese coast on fish welfare, employing a combination of qualitative and quantitative methods that included direct observation and analysis of vitality scores on deck and comparisons with circulating levels of physiological stress indicators. A major challenge in studying fish welfare during wild capture lies in how to assess and quantify welfare conditions while the fish remains underwater and in contact with the fishing gear. Assuming that the fishing event is bound to greatly decrease fish welfare, we decided to evaluate the variability of welfare indicators on arrival, as proxies for the impact of the fishing process, and the effects it may have, together with handling and environmental conditions, on further reduction of welfare whilst on board, considering that slaughter is not immediate in these operations. We evaluated the time-course of welfare decline from the time of arrival on deck and correlated with the handling and other conditions during storage on-board in four of the most common species targeted by this fishery, identifying which factors and indicators can contribute to or predict the decrease in welfare. Although ultimate goals must focus on improving the capture process itself, the primary objective is to provide accurate measurements of physiological and behavioral indicators occurring post-haul, related to fish welfare in the final stages of the fishing process, that can be used to evaluate the relevance and suggest mitigating solutions that can be practically implemented on-board the vessels of the small-scale fleet.

## 2. Materials and Methods

### 2.1. Study area

The study area comprised a section of coastal waters off southern mainland Portugal, also known as the Algarve (Figure 1). The Algarve coastline includes the south-west coast (∼50 km), from Odeceixe (37° 26’ N - 8°47’ W) to Cape São Vicente (37°1′ N - 8°59′ W), and the Southern coast (∼170 km extension), from Cape São Vicente to Vila Real de Santo António (37°11’ N - 7°25’ W). This coastal region has a very narrow continental shelf (5–20 km wide), with 2 major submarine canyons (S. Vicente and Portimão) and influenced locally by upwelling events, mostly occurring in the southwestern area. The southern area is also influenced by the more saline and warm waters of the Mediterranean Sea [37,38].

**Figure 1.**
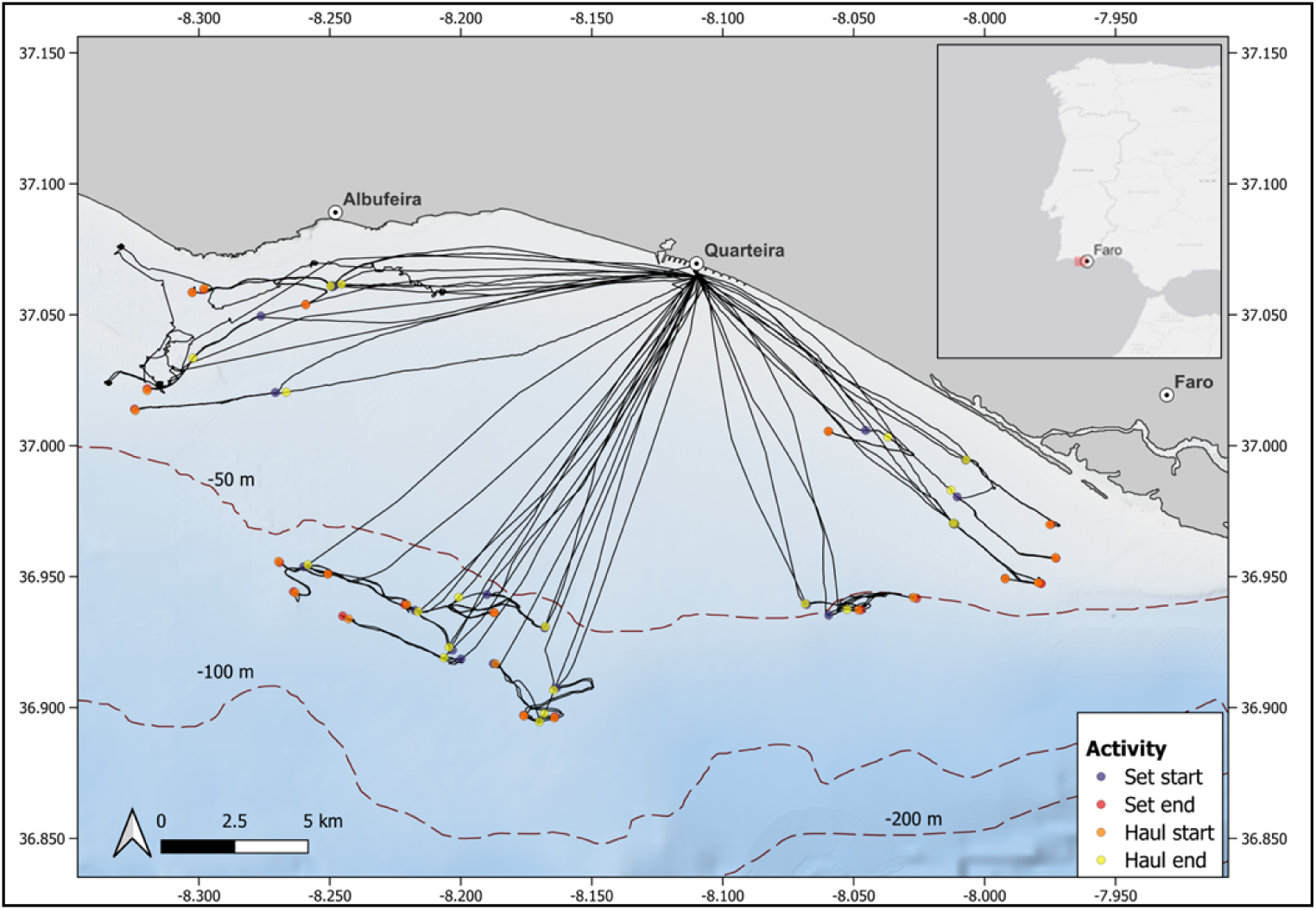
Map of the fishing tracks and fishing activities recorded from the sampling trips.

### 2.2. On-board sampling

Sampling took place from the port of Quarteira (37° 4’ N, 8° 6’ W) and the fishing operations were performed on the fishing grounds located off and towards east or west of the port (Figure 1). Overall, twenty-two sampling trips were undertaken but while vitality assessments were performed on all sampled trips, blood sampling for physiological analysis was carried out on 10 trips. The trips were performed on-board two commercial fishing vessels, one belonging to the small-scale fishing fleet, classified as local fleet (vessel size < 9 m) and one being a coastal vessel (vessel size > 9 m). Both operated bottom-set gill nets to target demersal fish species. The local vessel operated with a 60mm mesh size net that was 1.5m in height and 5000m long, while the coastal vessel operated with a mesh size of 78mm and a height and length of 3m and 4600m, respectively. The recorded depth of fishing ranged between 16m and 86m.

Trained observers went on-board the fishing vessels for opportunistic sampling on single-day trips that started typically before sunrise from the port of Quarteira. The fishing operation commenced on reaching the fishing grounds, and the time of net-setting and coordinates of the place, were recorded using a portable GPS device (GPSMAP^®^ 79, Garmin^®^). Time for setting the net usually lasted between 60-75 mins, post which, soaking time lasted around 60 mins before starting the hauling process. The hauling process spanned for about 150 minutes during which vitality assessments and blood sampling were performed on randomly selected specimens. Selection was conducted *ad libitum*, allowing fish to be hauled and disentangled by fishers while providing the observer with adequate space and time to make detailed observations.

The fish were selected separately in the sampling trips where we performed vitality assessments and the trips in which we sampled blood for physiological analyses. A. Vitality assessment: One of the samplers meticulously observed the hauling process. If a fish belonging to the desired species arrived, we requested the fisher to disentangle it and place it on our observation tray. We then completed the vitality assessment of that fish (until it reached vitality = 1) and repeated the process with another fish. B. Physiological analyses: When the fisher placed the fish on the tray we recorded the vitality and analysed vitality. The blood was sampled either at that stage (blue circles in Figure 2) or after reaching any other stage (green, yellow or orange circles in Figure 2). The cumulative stress in this case would be the effects of asphyxia, severe hypoxia, and desiccation as the fish struggle on the deck. Throughout the sampling process, we worked with the same fishers, thus maintaining constancy in the disentangling process.

**Figure 2.**
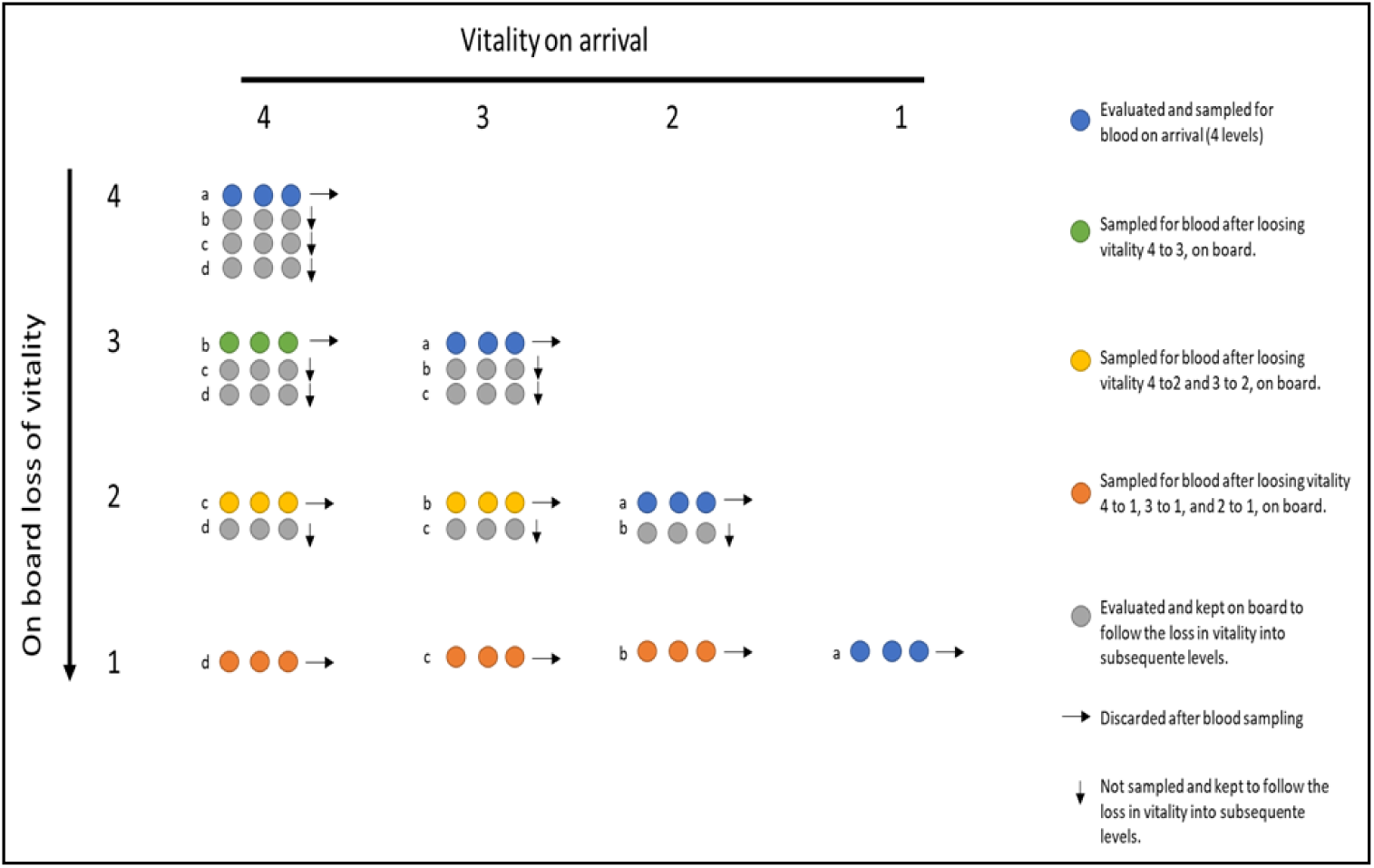
The matrix used for blood sampling. See legend for details.

Blood collection and fish manipulation were done under a FELASA type-C license issued by the DGAV (Portuguese Veterinarian Authority). All the fish manipulations were performed in accordance with the Portuguese legislation and the European Union guidelines (DL 113/2013, 2010/63/EU).

#### 2.2.1. Vitality assessments

Our experimental procedure aimed to simulate the current fishery process as close as possible. Vitality assessments were performed on four commercially important fish species that are commonly targeted by this gear, the striped red mullet (*Mullus surmuletus*, Family: Mullidae), the common pandora (*Pagellus erythrinus*, Family: Sparidae), the axillary seabream (*Pagellus acarne*, Family: Sparidae) and the two banded sea-bream (*Diplodus vulgaris*, Family: Sparidae) (hereafter, MS, PE, PA, and DV respectively). When a specimen belonging to any of those four species arrived on the deck, the observer recorded the time of landing, followed by measuring the total length of the specimen using a fish scale. The observers then registered the following variables while fish were in an observation tray:

— **Scale loss**: percentage of body surface without scales, considered as a proxy for wounds. Four categories were considered: 0-25%, 26-50%, 51-75% and 76-100 %.
— **Self-initiated movement**: fish jumping or otherwise moving by themselves; Indicator for vigor.
— **Reaction to touch**: fish initiates movement upon tail or body pinching; Indicator for maintenance of sensory capacity upon physical stimuli.
— **Opercular movement**: opening of the opercula; Indicator for maintenance of breathing reflexes.
— **Spasms**: Apparently random body jolting of shivering; Indicator for maintenance of basic physiological activity.
— **Eye roll / Vestibulo-Opercular Reflex (VOR)**: maintenance of the horizontal axis of the eyes when the fish is rotated. Indicator for conscious state [39].

Vitality was assessed every two minutes categorically recording their activity and reflexes against a pre-defined scale with four stages: 4. Highly Active (Flopping or other body movements on deck), 3. Less Active (Reaction to touch, regular opercular movement, spasms), 2. Lethargic (Random opercular movement, eye-roll/Vestibulo-Oculo reflex; VOR), and 1. Not responsive (No activity and responsiveness). However, it was observed during pilot trips that MS loses vitality rapidly, and thus, vitality was assessed over one-minute intervals in this species only. Handling was kept to a minimum to avoid conditioning the fish responses.

#### 2.2.2. Physiological analysis

Physiological stress indicators of captured fish were measured from blood samples collected according to the sampling matrix defined for this study (Figure 2). The matrix was established to relate the vitality of the individual at the moment it arrived on deck and the respective loss of vitality. This type of sampling was designed to compare not only the physiological status of the fish at the four vitality stages but also to evaluate the effect of the on-board procedures that lead to cumulative stress and negatively impact fish welfare. For instance, an individual with a given vitality score at arrival on deck (blue circles in Figure 2) was either sampled at that score or after losing vitality to any other lower score (green, yellow or orange circles in Figure 2). Physiological analyses were performed for the three species of sparids, PE, PA, and DV, but not for MS since most of the individuals of this species caught were either not responsive or lost vitality rapidly making blood collection unfeasible. Each fish was sampled for blood only once and removed from the matrix upon blood sampling, to avoid any confounding effects arising from this process.

Approximately 1 ml of blood was collected from the caudal vein with heparinised (1000 units/ml lithium heparin, Sigma-Aldrich^®^) 1ml-syringes (Terumo^®^) and needles (23G, 0.6X32 mm, Agani™, Terumo^®^) and immediately transferred to tubes containing 10 μl of heparin, mixed by agitation and stored in ice until the end of the trip. The samples were annotated with a two-digit code separated by a decimal. The digit before the decimal indicates the vitality at which the fish arrived on deck and the digit after the decimal indicates the vitality during blood sampling (Hence, a vitality stage of 4.2 suggests that a particular individual had arrived on the deck at vitality 4 and was sampled for blood at vitality 2).

### 2.3. Analytical techniques

Once in the laboratory, all blood samples were centrifuged at 9,000 g for 5 mins at 4°C (Heraeus Biofuge Stratos centrifuge) and the plasma was stored at -80°C until further analysis. The plasma levels of cortisol, glucose, lactate, and osmolality were measured to determine the extent of physiological stress on the captured fish. Plasma glucose and lactate concentrations were estimated with enzymatic-colorimetric methods adapted for 96-well plates using commercial kits (Spinreact ref.1001192 and ref.1001330, respectively). Final absorbances were measured at λ=505 nm using the Multi-mode Microplate reader Multiskan^®^ GO (Thermo Fisher Scientific Inc.) The results were expressed in mmol/L. Plasma osmolality was measured with a vapour pressure osmometer (Vapro^®^ Wescor 5520) and expressed in mOsm/Kg. Cortisol concentrations were estimated by radioimmunoassay (RIA) as previously described [39,40]. Briefly, the concentration of cortisol (F) was determined using a 1/200 or 1/400 dilution (in 0.01M PBS with gelatine, pH 7.6) of heat-denatured (1h at 80°C) plasma with a specific F antiserum (Anti-Cortisol 20 CR50, Fitzgerald^®^) and a radiotracer [1,2,6,7-3H] Cortisol (TRK407 Specific activity 50-90 Ci/mmol, Amersham Biosciences^®^). The samples, along with tritiated cortisol were incubated in a fixed quantity of antisera for at least 16 hours. The antibody-bound and free fractions were separated using charcoal, and a scintillation cocktail was added and thoroughly mixed. Radioactivity in the bound fraction was measured using a liquid scintillation analyzer (Tri-Crab 4810 TR^®^, Perkin-Elmer). The results were expressed in ng/ml.

### 2.4. Environmental data

The fishing depth was recorded using the fish finder devices present on-board the fishing vessels. Modelled data for Sea Surface Temperature (SST) and the temperature at the depth of fishing corresponding to the point and time of hauling for each fishing trip were obtained from products available in the Copernicus^®^ marine database (https://data.marine.copernicus.eu/). Likewise, data for atmospheric temperatures were obtained from the Copernicus^®^ climate store (https://cds.climate.copernicus.eu/) and Open-Meteo™ API (https://open-meteo.com/), respectively. The spatio-temporal resolution of the temperature variables is mentioned in table 1.

**Table 1:** List of fixed effect variables used in the Cumulative Linked Mixed Models (CLMMs) and Generalised Linear Models (GLMs)

| Category | Variable | Unit | Citation | Resolution |
| --- | --- | --- | --- | --- |
| Biological | Species | - | Present study |  |
|  | Length | Cm | [31] |  |
|  | Scale loss (Injuries) | % | [32] |  |
| Technical | Mesh Size | Mm | [32] |  |
|  | Depth of fishing | M | [45] |  |
| Environmental | Fishing depth temperature | °C | [45] | Temporal: Daily mean<br>Spatial: 0.083 X 0.083 deg |
|  | Sea surface temperature (SST) | °C | [28] | Temporal: Hourly mean<br>Spatial: 0.0625 X 0.0625 deg |
|  | Atmospheric Temperature | °C | Present study | Temporal: Hourly mean<br>Spatial: 0.28125 X 0.28125 deg |

### 2.5. Data analysis

All the subsequent data analyses were performed using the software R-Studio (stats ver. 4.3.0, R Core Team 2021) [42]. For the data obtained from the vitality assessments, the proportion of fish that landed on deck at the four vitality stages was calculated for each species. Non-parametric statistics were used when the data were non-normal and/or had heteroscedasticity. The loss of vitality post-hauling was assessed, initially, by comparing the time until on-board unresponsiveness between species with a Kruskal-Wallis test followed by a *post-hoc* analysis consisting of the Dunn’s test with a Bonferroni correction for pairwise comparisons. Furthermore, polynomial regressions were conducted to determine and compare between species, the rate of loss of vitality on the deck of the fishing vessel. Cumulative Linked-Mixed Models (CLMMs) fitted with the Laplace’s approximation were employed to enable the prediction of the vitality stage at arrival using the ‘clmm2’ function the package ‘ordinal’ [43]. The date of fishing was selected as the random factor to account for any variability that might have arisen as a result of several unaccounted factors during that fishing trip. Furthermore, to predict the duration of activity post-landing on the fishing deck, a Generalised Linear Model (GLM) with a negative-binomial function and log link was fit using the package ‘MASS’ [44]. Three categories of predictors were chosen, namely biological, technical, and environmental, based on past literature (Table 1).

The dependent variable (For CLMMs: The vitality stage at arrival on deck; For GLMs: Number of time intervals until non-responsiveness on deck) was modelled against all the non-correlated variables in a saturated model followed by a stepwise elimination process to select the most robust model. The best fit was chosen using the Akaike Information Criteria (AIC) and the principle of parsimony [47]. Modelling was done, first for all the species, and later, sub-models were fit for the individual species. It is imperative to note that the GLMs for PA and MS were fit using the Poisson function since it gave a better fit. To display the CLMM predictions graphically, the most robust model was fit using the ‘polr’ function in MASS without the random term, followed by a probabilistic plot. The plasma concentrations of the stress parameters between vitality stages were compared for each species using a single-way analysis of variances (ANOVA) or the non-parametric Kruskal-Wallis test when necessary. Since each species differs in the duration and magnitude of its stress response, between-species comparisons were deemed unsuitable. A significant cut-off was set at *p* < 0.05 and all the values are presented as mean ± s.e.m., unless otherwise stated. The figures were created using the R-Studio package “ggplot2” [48], while the map was created on the software Q-GIS Desktop (ver. 3.30.0).

## 3. Results

### 3.1. Vitality assessments

Vitality assessments were performed on 403 individual specimens belonging to the four n = 100species (DV: n = 100, avg. length = 21 ± 0.28 cm, avg. weight = 118.61 ± 6.79 g ; MS: n = 102, avg. length = 23.7 ± 0.26 cm, avg. weight = 181.92 ± 6.48 g PA: , avg. length = 22.4 ± 0.27 cm, avg. weight = 158.21 ± 6.65 g; PE: n = 101, avg. length = 22.6 ± 0.28 cm, avg. weight = 148.54 ± 6.84 g). In DV, most individuals were hauled on deck with the highest vitality scores of 4 and 3 (Figure 3a). In contrast, almost 50% of the mullids (MS) arrived on the deck inactive/ unresponsive (vitality = 1) or lethargic (vitality = 2) and became unresponsive within the first minutes (Figure 3b). In the other two sparids PA and PE, individuals showed similar trends to DV, with most fish landing on deck at a vitality score of 4 (Figure 3c and 3d).

**Figure 3.**
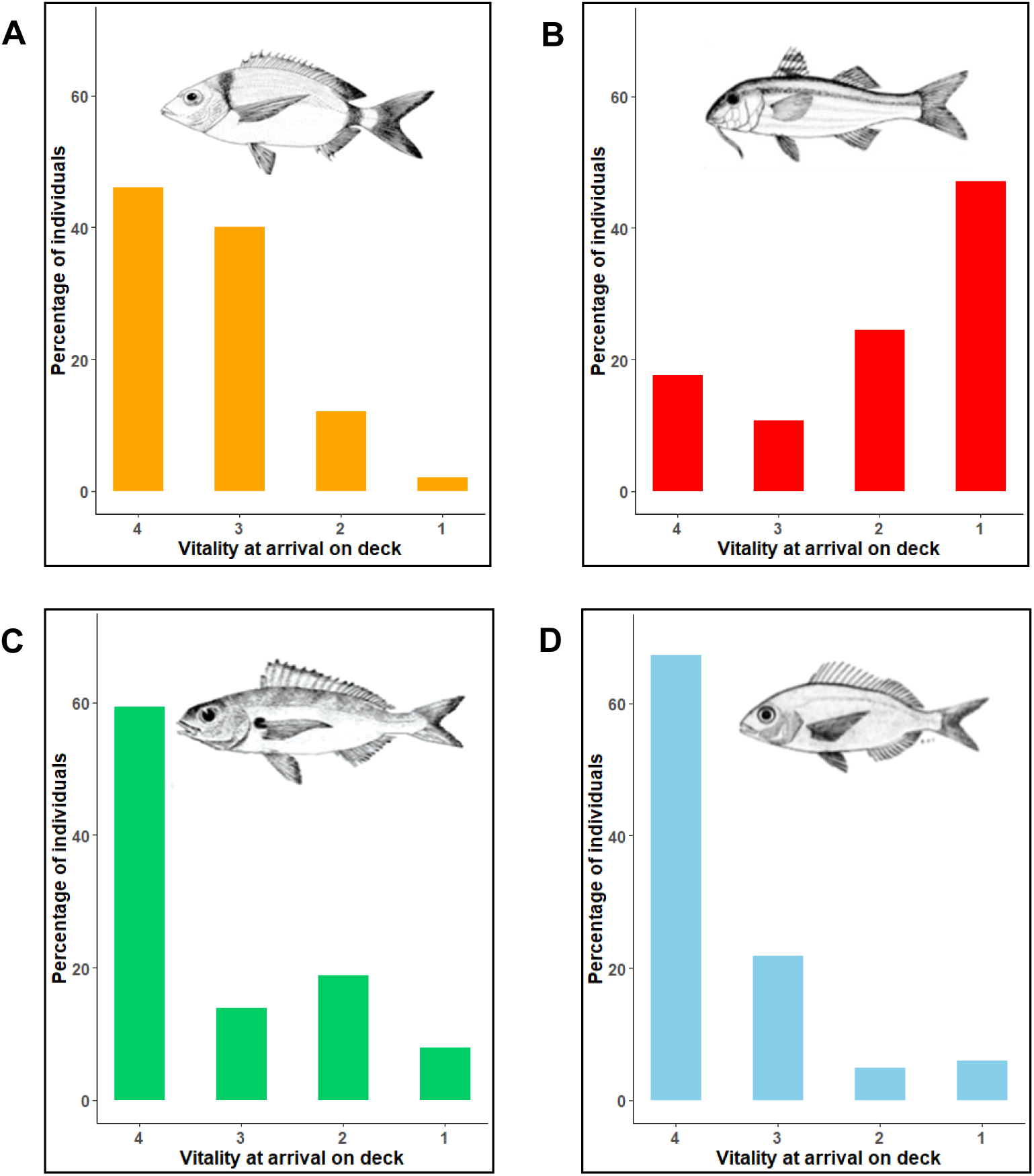
The proportion of fish landed on deck at the four vitality stages. (a) DV: Two-banded seabream (*Diplodus vulgaris*); (b) MS: Red mullet (*Mullus surmuletus*); (c) PA: Axillary seabream (*Pagellus acarne*); (d) PE: Common pandora (*Pagellus erythrinus*). (Vitality scale: 4 = highly active, 3 = less active, 2 = lethargic and 1 = unresponsive.).

The duration of activity on the deck of the fishing vessel (χ² = 80.49, p = 0, Kruskal-Wallis test), with the Dunn’s test, revealed significant differences between all the pairs except DV and PE (Figure 4a). DV was shown to have the slowest loss of vitality (Figure 4b1) even though it had the lowest average vitality when landed on deck amongst sparids, with one individual responsive for as long as 82 minutes after we started examining it. In contrast, MS showed the most rapid loss of vitality with the maximum time on deck being 4 minutes (Figure 4b2). Likewise, PA lost vitality most rapidly between sparids along with an average vitality at the time of arrival on deck being around 3 (Figure 4b3). PE had the highest average vitality at the time of landing on the vessel (Figure 4b4), but it lost vitality at a relatively slower rate when compared to PA.

**Figure 4.**
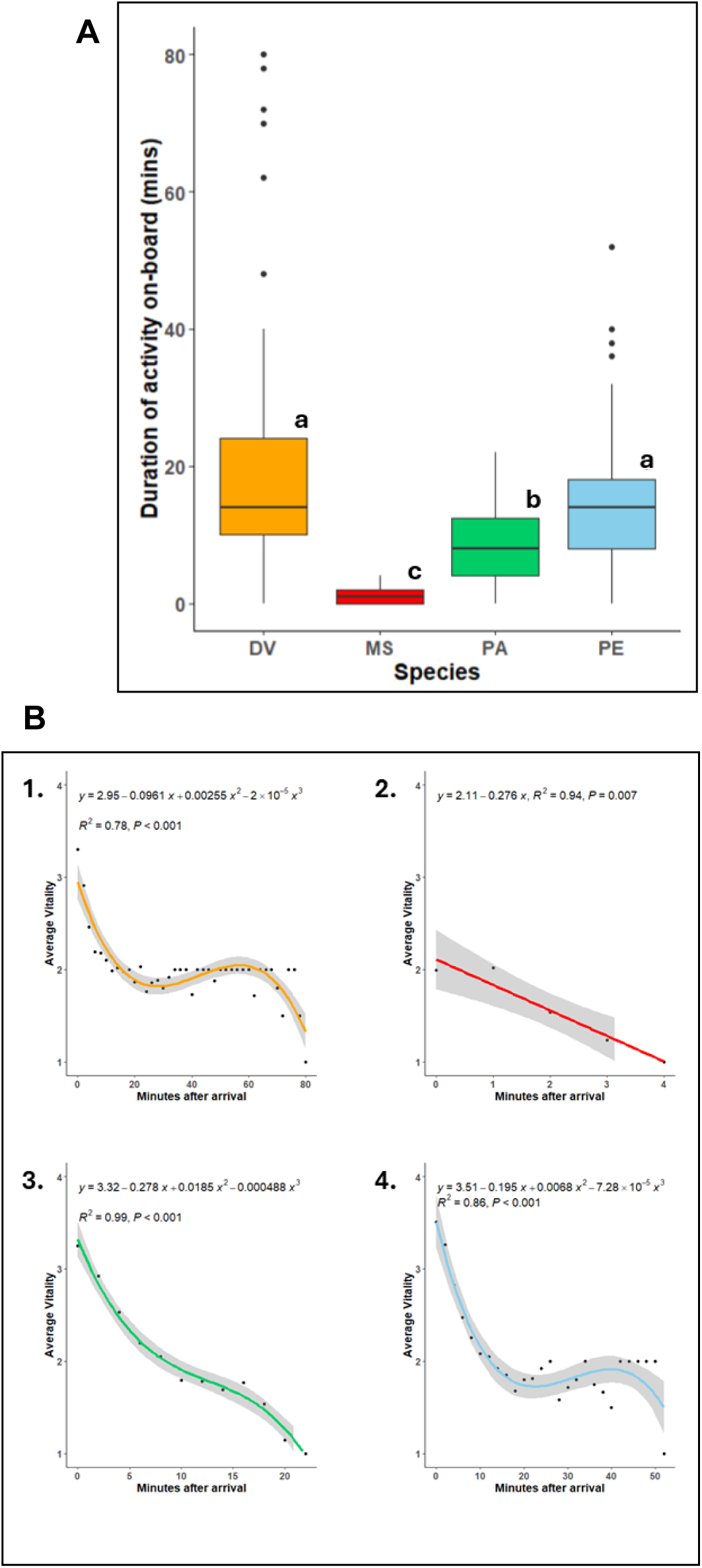
(a) A comparison of the time until on-board unresponsiveness between species. Statistical differences (*p* < 0.05) are signaled with different letters; (b) Polynomial regressions showcasing the average loss of vitality on-board concerning time. (Each species is indicated by a different colour 1. DV: Two-banded seabream (*Diplodus vulgaris*), 2. MS: Red mullet (*Mullus surmuletus*), 3. PA: Axillary seabream (*Pagellus acarne*), 4. PE: Common pandora (*Pagellus erythrinus*)). (Vitality scale: 4 = highly active, 3 = less active, 2 = lethargic, and 1 = unresponsive. Time period refers to observations at every minute for red mullet and every 2 minutes for other species, grey shades: 95% confidence intervals; black dots: average vitality at a particular time).

CLMMs indicated that the vitality of fish at arrival was largely a function of two variables: species (Figure 5a) and the degree of scale loss (Figure 5b). As expected, MS had the highest probability of arriving at the lower vitality scores of 1 and 2 (*p* < 0.001). Contrastingly, the sparids had a greater probability of being landed on deck at higher vitality scores of 3 and 4, with PE individuals generally predicted to be the most vital at the time of landing. Likewise, the degree of scale loss (Figure 5b) was predicted to have a detrimental effect on vitality. This implies that individuals with a given scale loss rating of 2 (26 – 50% scale loss) had a significantly lower probability of being highly vital at the time of landing on the fishing vessel (*p* = 0.003). This prediction, however, should be interpreted prudently as most fish being sampled had minimal or no scale loss (scale loss rating = 1, n = 370), as compared to those with a scale loss rating of 2 (n = 33). Moreover, scale loss was observed only in DV and MS and a glance at the sub-models fit for individual species shows similar predictions concerning scale loss for both species (Supplementary table S1).

**Figure 5.**
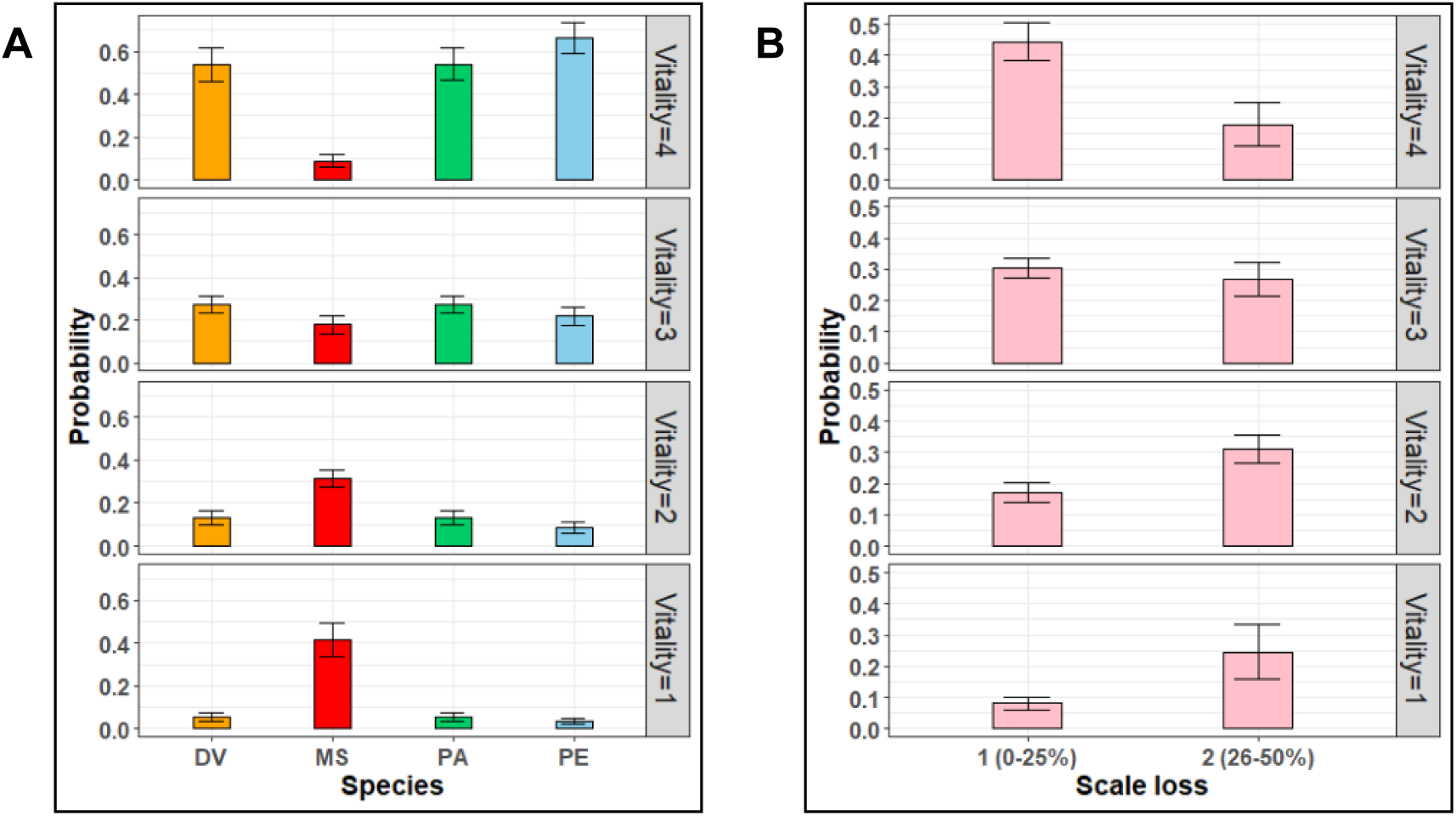
The probabilistic plots showcasing the probability of fish landing at a particular vitality stage based on the CLMM model predictions. (a) Species: DV: Two-banded seabream (*Diplodus vulgaris*), MS: Red mullet (*Mullus surmuletus*), PA: Axillary seabream (*Pagellus acarne*), PE: Common pandora (*Pagellus erythrinus*); (b) Scale loss in DV and MS, 1: 0 – 25% scale loss on the body, 2: 26-50% scale loss on the body. (Vitality scale: 4 = highly active, 3 = less active, 2 = lethargic; and 1 = unresponsive. Error bars: ± standard error.).

Amongst the GLMs that were fit to predict the duration of activity on-board the vessel due to various factors, the best-selected model was the one having species, length, scale loss, fishing depth temperature, sea surface temperature, and atmospheric temperature as the significant predictors (Table 2).

**Table 2.**
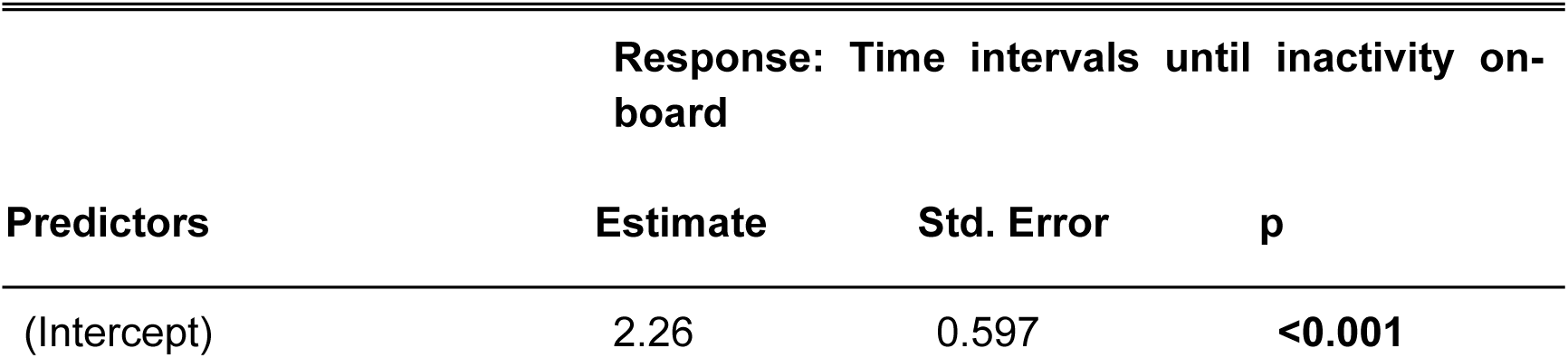
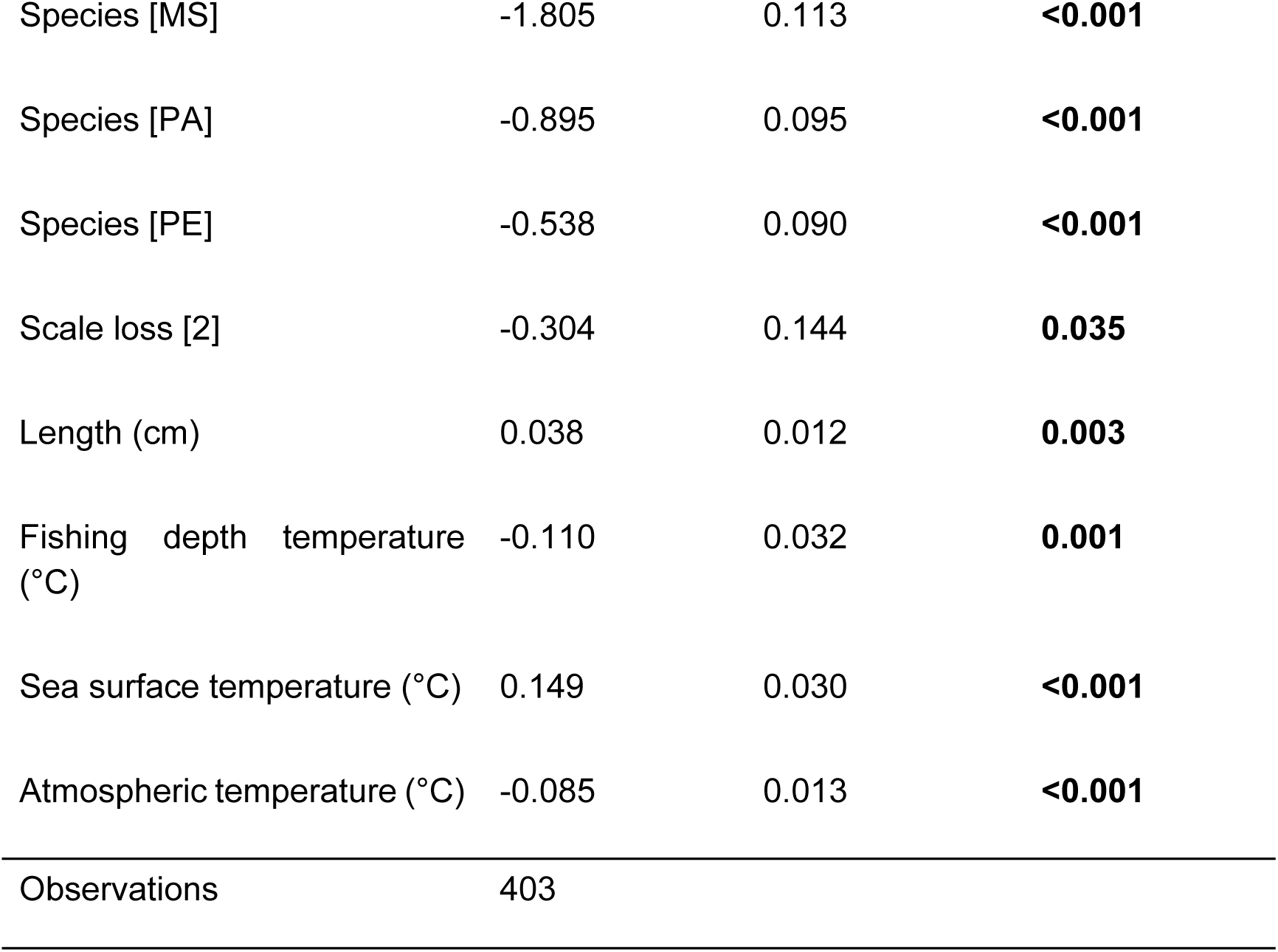
The Estimate, standard error, and the p- values (p: Pr(>|z|)) derived from the negative-binomial GLM that was fitted to predict the impact of several biological, operational, and environmental predictors on the duration of fish activity on-board the fishing vessel.

| Predictors | Response: Time intervals until inactivity on-board |  |  |
| --- | --- | --- | --- |
|  | Estimate | Std. Error | p |
| (Intercept) | 2.26 | 0.597 | <0.001 |
| Species [MS] | -1.805 | 0.113 | **<0.001** |
| Species [PA] | -0.895 | 0.095 | **<0.001** |
| Species [PE] | -0.538 | 0.090 | **<0.001** |
| Scale loss [2] | -0.304 | 0.144 | **0.035** |
| Length (cm) | 0.038 | 0.012 | **0.003** |
| Fishing depth temperature (°C) | -0.110 | 0.032 | **0.001** |
| Sea surface temperature (°C) | 0.149 | 0.030 | **<0.001** |
| Atmospheric temperature (°C) | -0.085 | 0.013 | **<0.001** |
| Observations | 403 |  |  |

The model predicted a significant difference in the duration of activity between species. Likewise, 26 – 50% scale loss (scale loss rating: 2) significantly reduced the duration of activity (z = -2.109, *p* = 0.035). The model predictions also showed a significant increase in the duration of activity with increasing fish length (z = 2.962, *p* = 0.003), which indicates that large fish may struggle more and take longer to become unresponsive while on deck compared to smaller ones.

Amongst the environmental variables, atmospheric temperature (z = -6.241, *p* < 0.001) influenced the duration of activity with a sharp decrease of activity with increasing temperatures. A similar effect was observed in the case of temperature at the depth of fishing (z = -4.862, *p* = 0.001). However, an opposite trend was noticed in the case of sea surface temperatures (z = 2.58, *p* < 0.001), wherein the duration of activity was positively related to the sea surface temperature. Importantly, the sub-model for DV predicted a significant decrease (z = 5.5, *p* < 0.001) in the duration of activity with increasing depth in DV (Figure 6a), indicating the effect of barotrauma.

**Figure 6.**
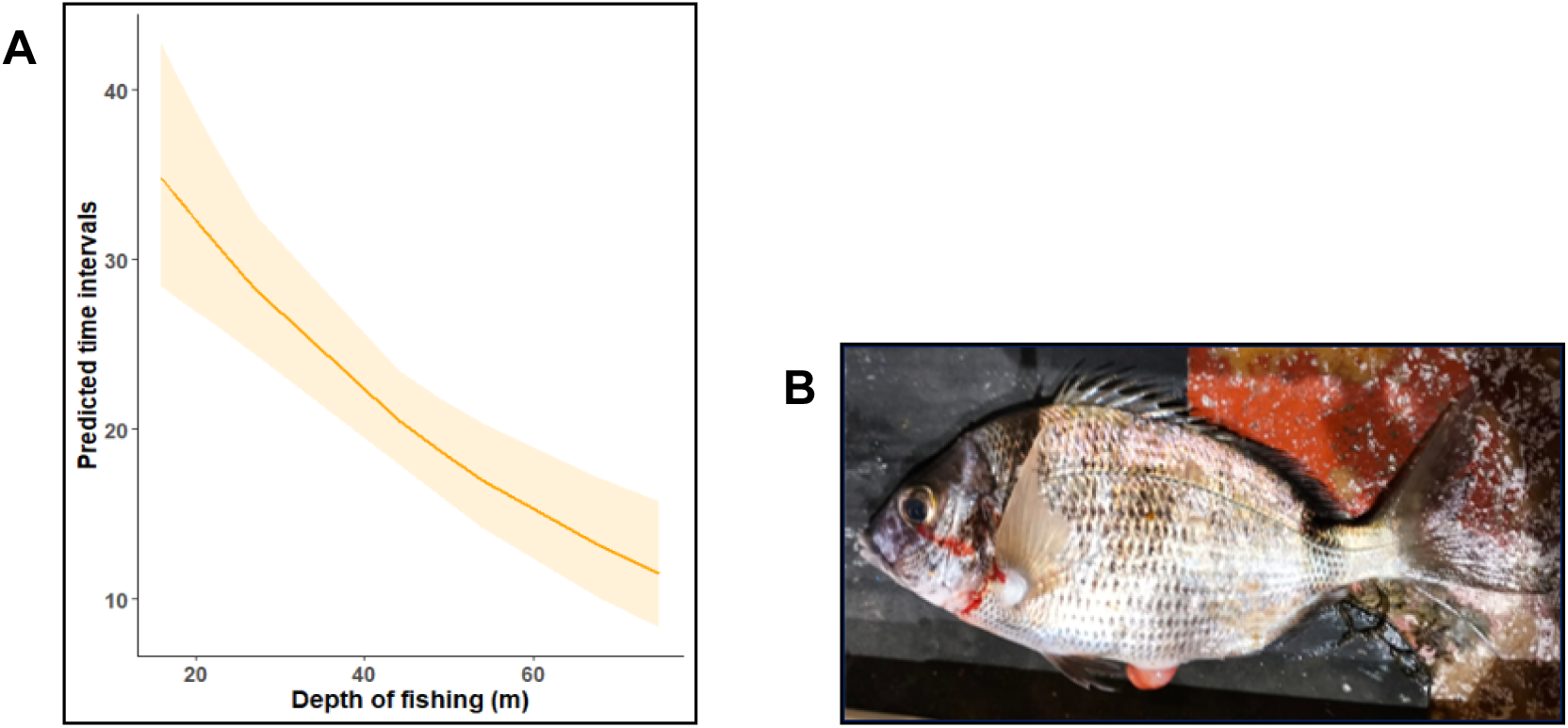
(a) GLM prediction of the response variable - time intervals until inactivity and unresponsiveness as a function of the explanatory variable-depth of fishing (m) (Orange shading: 95% confidence intervals); (b) Evidence for signs of barotrauma in DV as suggested by the protrusion of the swim bladder (Picture Credit: Magda Frade).

### 3.2. Physiological analysis

Physiological analyses were performed on a total of 183 fish (59 DVs, 64 PAs, and 60 PEs, respectively). Average levels of cortisol (Figure 7a) were observed to vary non-significantly (Kruskal-Wallis test: *p* = 0.50) between the vitality stages in DV and PA. In PE, the highest average concentration of cortisol was observed in the lowest vitality stage of 1.1 (135.44 ± 53.30 ng/ml), whereas the higher vitality stage of 4.3 had the lowest cortisol concentration (25.14 ± 12.59 ng/ml). However, no statistically significant differences were found between vitality stages (Kruskal-Wallis test: *p* = 0.50).

**Figure 7:**
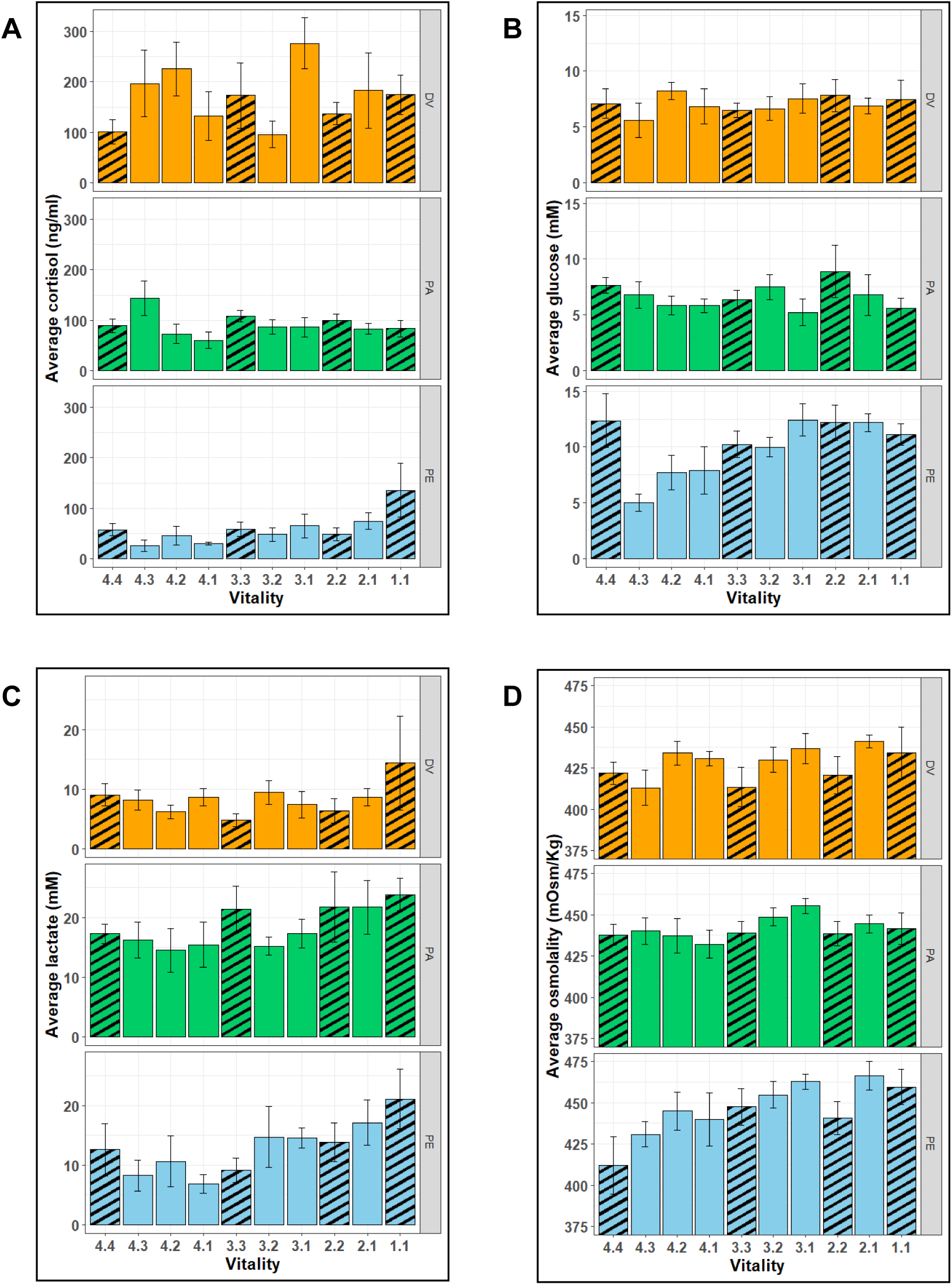
A comparison of average levels of the plasma parameters measured in Sparids at different vitality stages. (a) cortisol (ng/ml); (b) glucose (mM); (c) lactate (mM); (d) osmolality (mOsm/Kg). (Species: DV: Two-banded seabream (*Diplodus vulgaris*), PA: Axillary seabream (*Pagellus acarne*); PE: Common pandora (*Pagellus erythrinus*)). (Vitality scale: 4 = highly active, 3 = less active, 2 = lethargic and 1 = unresponsive. Error bars: ± standard error. Vitality stage: The digit before the decimal indicates the vitality at which the fish arrived on deck and the digit after the decimal indicates the vitality at the time of blood sample, i.e. a vitality stage of 4.2 suggests a fish that was landed at vitality 4 and was sampled at vitality 2. Striped bars represent vitality stages wherein the vitality at arrival on deck were the same as the vitality at the rime of sampling. The results of the statistical significance tests can be found in Supplementary tables S3 and S4).

The variations in the average plasma concentration of glucose (Figure 7b) between vitality stages were not significant in DV and PA (Anova: *p* = 0.38 for DV; *p* = 0.48 for PA). However, considerable differences (Anova: *p* < 0.001) in glucose concentrations were found in PE, with the levels at stage 4.3 (5.02 ± 0.76 mM) being significantly lower (*p* < 0.05) than those at stages 4.4 (12.36 ± 2.41 mM), 3.1 (12.43 ± 1.44 mM), 2.2 (12.19 ± 1.55 mM), and 2.1 (12.19 ± 0.80 mM).

Lactate concentrations (Figure 7c) did not differ significantly between vitality stages for any of the three species (Anova: *p* = 0.48 for DV; Kruskal-Wallis test: *p* = 0.44 for PA; *p* = 0.11 for PE). However, the highest average plasma concentrations in all the three sparids were noted for the lowest vitality stage, i.e. 1.1 (14.39 ± 7.89 mM in DV; 23.85 ± 2.66 mM in PA; 21.11 ± 5.01 mM in PE). Likewise, osmolality levels (Figure 7d) did not vary significantly across vitality stages in either DV or PA (Anova, *p* = 0.35 for DV; *p* = 0.71 for PA), but significant differences were observed in PE (Anova, *p* < 0.05), in which the osmolality of fish sampled at a comparatively lower stage 2.1 (466.33 ± 8.33 mOsm/Kg) was significantly higher (*p* < 0.05) than the ones sampled at the highest vitality stage 4.4 (412 ± 17.58 mOsm/Kg).

## 4. Discussion

This study integrated behavioural and physiological stress indicators to provide new insights into the welfare state of fish caught in bottom-set gill net fisheries. Results showed species-specific differences in vitality at capture and its decline on deck. Scale loss was identified as a key factor affecting vitality. Environmental conditions and fish size were also significant predictors of vitality loss. Physiological assessments confirmed high stress levels, with cortisol elevated across all vitality stages and increased lactate and osmolality in fish with the lowest vitality, and/or longer periods on deck, indicating further loss of welfare after capture. This is one of the first studies attempting to measure welfare indicators in wild-capture fish at hauling and during subsequent onboard operations and emphasises the importance of improved fishing and handling practices to minimise negative welfare impacts in set-net fisheries.

### 4.1. Vitality assessment

It is well known that fish species differ significantly in their ability to tolerate stress [49]. Our results indicate that the mullid, *M. surmuletus* was less resistant to the stress and physical demands of the fishing operation and hence, go unresponsive faster compared to the sparids, which, assuming similar soak times, had comparatively greater tolerance to capture and post-capture stress. Between the three sparids, PE can be assumed to be the most stress-tolerant to the stressors of capture with most individuals being landed on deck at the highest vitality score of 4. However, it must be stressed that the current approach could not determine when fish first contacted the gear and therefore soaking time could not be used as a factor in our analysis, despite its undoubted importance.

All fish were observed to experience multiple stressors between the time they are hauled and their eventual loss of activity and responsiveness on deck. They included physical injury caused by fisher activities like disentangling the fish from the net and throwing them in a storage tray or bucket [50,51], followed by asphyxiation, hypoxia, heating, fatigue, and dehydration which eventually lead to the death of the fish [9,52]. The duration of activity and the loss of vitality varied significantly between the four species, with DV and PE showing similar trends. Even though DV had a low average vitality at the time of landing, it had a slower rate of vitality loss and took a long time to go inactive. Therefore, DV not only had the highest tolerance to the on-board stressors amongst the four species but was also the most tolerant to the cumulative stress from all the stressors. However, higher stress tolerance does not imply better welfare of that species. Contrarily, being stressed for a longer duration suggests that the fish have to struggle, and potentially, suffer more [53]. Several specimens of DV showed muscular stiffening before death (data not shown). The exhaustion during capture followed by a long-duration asphyxiation period on deck might have contributed to a scarcity of Adenosine Tri-Phosphate (ATP), leading to the onset of *rigor mortis*. This observation is in conjunction with several studies that showcase an early onset of rigor in fish that experienced pre-mortality capture and handling stress, especially at higher temperatures [35,54,55].

One of the main criteria for poor welfare is exposure to stress for long durations of time [36] and it is well known that the vitality of a fish is negatively correlated to the amount of stress it experiences [56]. Since vitality and welfare are positively correlated [25], it is reasonable to assume that a fish struggling for a longer duration on-board dies with poorer welfare as compared to the one which loses vitality rapidly and experiences a quick death. Based solely on the vitality assessments, this study suggests that DV experienced the worst welfare at the time of perceived death amongst the four species because it struggled for a long duration on deck in a lethargic condition (vitality 2) witnessed by the lack of body movement, near-complete reflex impairment, irregular opercular movement, and infrequent eye-roll reflex before its eventual inactivity. Thus, it is proposed that while assessing welfare in fish, the pre-inactive vitality condition should also be considered along with the duration of activity post-capture [57].

Various biological, operational, and environmental variables have been used in literature to explain the amount of stress experienced by fish during fishing operations [31,32,46]. Scale loss can be considered as an important factor since it not only predicted the reduction of the vitality of fish at arrival, but also significantly shortened the duration of activity in both the species it was observed in (i.e. MS and DV). Scale loss and other type of external injuries have been found to be one of the most important factors that compromise fish welfare [26]. They can severely impact the welfare of the fish being caught by causing a physiological stress response [59–61], osmoregulatory impairment [62,63], blood loss, inflammation, and higher susceptibility to infections amongst others [64]. Such external injuries have been consistently noted to have increased the proportion of discard mortalities amongst fishes that experience fishing stress [19,65,66]. With regards to this study, it can be expected that scale loss elevated the stress levels in fish by exacerbating the osmoregulatory distress when the fish struggled after being caught in the net and dehydration after being landed on the vessel.

Several studies have shown smaller fish having less tolerance to capture and post-capture stressors and thus, had a higher behavioral impairment, earlier mortalities, and lower percentages of discard survival [11,19,58,67,68]. This was predicted by our full GLM model, wherein fish length significantly increased the activity duration. Contrary to this, a study conducted on the discard mortality of red mullet (*Mullus barbatus*) due to the stressors of bottom trawling resulted into the larger fish having a higher probability of mortality [69]. This is in conjunction with the predictions of our sub-model for MS, which predicted bigger fish to generally arrive on deck at lower vitalities. The net mesh size, an operational variable, was not significant in the final model consisting of all the species and was seen to reduce its robustness. However, it showed a large impact in the sub-model fit for PA (Supplementary tables S1, S2, showing species-specific models), with individual fish landed on deck at higher vitalities and displaying longer activity times when caught using a mesh size of 78mm, when compared to a relatively smaller mesh size of 60mm. The rate of ascent, or the hauling rate is also linked to the depth of fishing, a variable that was significant in the case of DV. Fish that are fished from greater depths and hauled at a faster ascent rate experience higher stress as a result of barotrauma [28], and hence, attain inactivity at a relatively faster rate. Again, soaking time could not be considered. However, we worked under the assumption that in 22 trips, the soak time of the 403 fish would have distributed randomly throughout the fishing period.

We found high atmospheric temperature to reduce the time it takes for an individual to completely lose its vitality on the fishing vessel. This could be due to the fact that elevated atmospheric temperature acts as synergistic stressor to fish that are already experiencing extreme stress due to hypoxia by increasing metabolic rates [70], and oxygen demands [71,72], which could not be fulfilled. Similar trends were obtained for temperature at the depth of fishing in the final model as well as in the sub-models for MS and PA, with increasing temperatures also predicting a reduced vitality at arrival for MS and PE. These results are in corroboration with several studies that have observed an increase in physiological stress and mortality with elevated temperatures during the capture process [31,45,67,73]. Lastly, water temperature (measured as SST) showed a positive relation with the duration of activity. Elevated SSTs were also predicted to be responsible for increasing the vitality at landing in DV and PA. Elevated SSTs implied reduced temperature difference between the surface and the atmospheric temperature, hence reducing the temperature shock experienced by fish when they are landed on deck.

### 4.2. Physiological analyses

The present study focused on determining if the levels of stress parameters vary between the four vitality scores in the targeted species as well the effects of cumulative fishing stressors on those parameters. To the best of our knowledge, is the first one to have measured cortisol in the sparid species, the DV and the PA, and one of the few with similar data on the PE [74]. A previous study determined the resting level of plasma cortisol in sparid fishes to be lying in the range of 1 ng/ml to 10 ng/ml, with elevated levels of 90 – 100 ng/ml measured from fish experiencing acute stress [75]. Likewise, another one reported reference cortisol concentrations in the sparids *Lithognathus mormyrus* (17.87 ± 3.17 ng/ml), *Dentex dentex* (15.20 ± 3.62 ng/ml), and *Sparus aurata* (4.61 ± 0.88 ng/ml) acclimatised under optimal conditions with minimal stress [76]. Whereas, a study performed on blackspot seabreams (*Pagellus bogovareo*) that had experienced stress in the course of hook and line fishing had elevated plasma cortisol levels of almost 100 ng/ml at the time of landing on-board [77]. Likewise, in PE, one of our target species, exposed to stress due to chasing followed by air exposure showed maximum average plasma concentrations of approximately 300 ng/ml an hour after exposure to the stressors [74]. We report markedly high average cortisol levels in fish at the time of landing on deck (DV: 174.40 ± 39.53 ng/ml at vitality 1; PA: 108.18 ± 11.87 ng/ml at vitality 3; PE: 135.44 ± 53.30 ng/ml at vitality 1) as well as in fish that had experienced cumulative stress on-board the vessel (DV: 276.13 ± 50.69 ng/ml at vitality stage 3.1; PA: 143.94 ± 34.01 ng/ml at vitality stage 4.3). We had presumed the cortisol levels to rise with decreasing vitality as a reduced vitality is concurrent with increased stress. However, no such effect was observed for DV and PA, even in those individuals tested for the build-up of cumulative stress due to the on-board activities. In the case of PE, fish that arrived inactive (vitality stage = 1.1) had the highest average plasma concentration of cortisol. The lack of an observable pattern of increasing cortisol was rather unexpected since air exposure is known to acutely increase blood cortisol in fish [78,79]. However, it must be considered that the fish in this study may had been struggling in the net for possibly 15 to 120 min and hence, the cortisol response had already been initiated or even plateaued. This is accentuated by the high levels of cortisol measured in fish when they arrived on deck , as compared to the control values of 1-21 ng/ml described for the sparids [75,80], but also contrasting to those reported recently in *Diplodus annularis* caught in trammel net fisheries (22.0 ± 5.3 ng/ml) [29]. Such dissimilarity (a 10-fold difference) is pondering, and may be related with species, detection methods, or reflect differential impacts between the fishing methodology used in the two studies. The plasma glucose concentrations were not significantly different between the four stages in the three species. The blood concentration of glucose depends on the level of glycogen reserves in the tissue, which varies between species and between individuals [81,82]. Unlike the simulated fishing experiments that are performed under controlled conditions [83], we performed the plasma analyses on samples obtained from wild fish in which the reserves of glycogen and thus, the concentration of blood glucose would be a function of the quantity and the quality of food consumed by individual fish before capture along with the duration and intensity of muscular exercise performed before and during capture.

Fish that landed on deck tended to show higher plasma lactate levels at the lower vitality stages in the three studied species. It is possible that the fish once gilled, wedged, snagged or entangled would struggle with impaired gill function to uptake oxygen and severe muscular exercise eventually leading to exhaustion. Thus, the following hypoxia would cause a build-up of lactate in the muscle cells and the blood. Similar results were reported in the striped bass, *Morone saxatilis* caught using gill nets [84]. This result is in agreement with a recent study performed on Atlantic mackerel *Scomber scombrus*, which observed a negative correlation between blood lactate concentration and vitality in fish that had experienced purse seining stressors [25]. Surprisingly, there was no significant rise in lactate concentrations while on-board, which was to be expected because of asphyxia-induced hypoxia and anaerobiotic muscular exercise during struggling, as indicated by various studies assessing the impact of air exposure on discard mortality [77,79,85].

Plasma osmolality showed an increasing trend with decreasing vitality in DV and PE. There was also some evidence of higher osmolality due to on-board stress with vitality stage 2.1 having the highest average osmolality in DV and PE, and stage 3.1 in PA. Air exposure on-board will likely lead to desiccation and water loss from the blood to muscles, increasing osmolality. It can also be presumed that the acute stress will lead to a quick secretion of catecholamines in the blood, which increasing gill permeability and the uptake of ions while still in seawater [86].

### 4.3. Study limitations

A major limitation of this analysis of vitality assessments is that the time at which a fish is caught in the net is not known. Therefore, there is no definite estimate of the time for which the fish was struggling in the net during the soaking time, which would have inevitably affected the vitality of each fish analysed. In general, longer soak times are said to be more detrimental to the welfare of fish and studies in the past have used the maximum soak time as a predictor for post-discard mortality [4,46]. However, within the design of this study, considering soaking time as a variable would have been misleading since it was not possible to collect individual soaking times for each fish. This means a fish caught during the beginning of the net set would have struggled and thus lost more vitality than the one caught at the end of the net set, but they would still be modelled against the same soak time.

The limitation of not knowing the exact time of capture is also relevant in this case because the stress experienced between the times of capture and landing can considerably affect the level of the four physiological parameters. Lastly, a drawback was the lack of data on the baseline levels of these parameters for the 3 species studied to compare our results with. The current study only compared the extent of physiological stress between the four vitality stages. However, comparing the levels obtained from these stressed fish to those in completely non-stressed fish is imperative to determine the true extent of stress experienced by these fish throughout the process of gillnet fishing.

## 5. Conclusions

Initiatives promoting fish welfare policies in fisheries are mounting, mostly based on evidence from the aquaculture industry. However, there is a lack of scientific information from wild fisheries operations that could be used to mitigate negative impacts to fish, promote better fishing standards that add value to the quality of the fish and support decision-making in fisheries management._–_This study assessed fish welfare post-haul in small-scale bottom-set gillnet fisheries using vitality assessments and physiological indicators. Sparids demonstrated greater resistance to fishing stressors than mullids, with DV showing the longest struggle under impaired reflexes. Multivariate analysis identified biological, operational, and environmental factors influencing welfare, with atmospheric temperature being the most critical variable in reducing responsiveness duration. Elevated cortisol levels across all species and vitality stages confirmed high stress, while increased lactate and osmolality indicated additional stress from on-board handling. Given the poor welfare conditions observed, mitigation measures such as reducing fishing depth or hauling speed, could be applied. Similarly, slight changes on post-capture process could benefit welfare. Fish often receive callous treatment once on deck and improvements on fish handling can be achieved using ramps leading to the storage units. Additionally, the implementation of immediate and humane slaughter methods are recommended to reduce further welfare loss, preserve quality and increase product value. However, challenges like vessel size, costs, lack of incentives, and limited awareness hinder their adoption and are a challenge, mainly for small-scale fisheries.

## Supplementary Materials

The following supporting information can be downloaded at: https://www.mdpi.com/xxx/s1, Table S1: The Estimate, standard error, and the p-values (*p*: Pr(>|z|)) derived from the Generalised linear models (GLMs) that were fit to predict the impact of several biological, operational, and environmental predictors on the duration of fish activity on the deck of the fishing vessel. (DV: Two-banded seabream (*Diplodus vulgaris*), MS: Red mullet (*Mullus surmuletus*), PA: Axillary seabream (*Pagellus acarne*), PE: Common pandora (*Pagellus erythrinus*); Table S2: The Odds ratios, CI: Confidence intervals, and the p-values derived from the Cumulative linked mixed models (CLMMs) that were fit to predict the impact of several biological, operational, and environmental predictors on the vitality at the time of landing on deck. (1. All species: The full model consisting of all the species, DV: Two-banded seabream (*Diplodus vulgaris*), MS: Red mullet (*Mullus surmuletus*), PA: Axillary seabream (*Pagellus acarne*); Vitality scale: 4 = highly active, 3 = less active, 2 = lethargic and 1 = unresponsive); Table S3: The p-values obtained from statistical significance tests performed to compare the levels of physiological stress parameters between vitality stages depending upon the vitality at arrival (DV: Two-banded seabream (Diplodus vulgaris), PA: Axillary seabream (Pagellus acarne), PE: Pagellus erythrinus; Anova: Analysis of variance, KW: Kruskal-Wallis test, t-test: Welch two sample t-test); Table S4: The p-values obtained from statistical significance tests performed to compare the levels of physiological stress parameters in vitality stages where the vitality at arrival was the same as vitality at sampling (stages 4.4, 3.3, 2.2, and 1.1) as indicated by the striped bars in Figure 7. (DV: Two-banded seabream (Diplodus vulgaris), PA: Axillary seabream (Pagellus acarne), PE: Pagellus erythrinus; Anova: Analysis of variance, KW: Kruskal-Wallis test).

## Abbreviations

The following abbreviations are used in this manuscript:

CVA: Categorical vitality assessment
GPS: Global positioning system
FELASA: Federation of European Laboratory Animal Science Associations
DGAV: Direção-Geral da Alimentação e Veterinária
MS: *Mullus surmuletus*
PE: *Pagellus erythrinus*
PA: *Pagellus acarne*
DV: *Dilpodus vulgaris*
VOR: Vestibulo-oculo reflex
RIA: Radioimmunoassay
SST: Sea surface temperature
CLMM: Cumulative linked-mixed model
AIC: Akaike information criteria
ANOVA: Analysis of variance
GIS: Geographic Information system
ATP: Adenosine tri-phosphate

## Author Contributions

Conceptualization, P.G., J.G., J.S. and A.M.; methodology, P.G., J.G., J.S., R.C. and A.M.; formal analysis, V.S. and L.B.; investigation, A.M., V.S., M.F. and R.C.; data curation, V.S., M.F., L.B. and R.C.; writing—original draft preparation, V.S. and R.C.; writing and reviewing— P.G., A.M., J.G., J.S.; visualization, V.S.; supervision, P.G., A.M., J.G.; project administration, P.G., J.G. and J.S.; funding acquisition, J.S., P.G. and J.G.. All authors have read and agreed to the published version of the manuscript.

## Acknowledgments

The authors are grateful to the skipper and the crew of the fishing vessels Zé Rita and Comendador for their active support and cooperation. We thank the other partners of Carefish/catch consortium, *fair-fish international*, Friends of the Sea and Demos for support and fruitful discussions.

## Institutional Review Board Statement

The observations were conducted aboard commercial fishing vessels and fish handling was performed by fishermen in their routine for collecting and storing the fish. Procedures for vitality scoring and blood sampling were performed under compliance with internal ethics boards and the directives issued by Direção Geral de Alimentação e Veterinária (DGAV, Licence reference 0421/000/000/2021), Ministério da Agricultura, Florestas e Desenvolvimento Rural, Portugal in compliance with the European (Directive 2010/63/EU) and Portuguese legislation (Decreto-Lei no. 113/2013 de 7 de Agosto) for the use of laboratory animals. All procedures were conducted or supervised by trained scientists under Group-C licenses issued by the DGAV, Ministério da Agricultura, Florestas e Desenvolvimento Rural, Portugal.

## Data Availability Statement

Data will currently be made available on request. A complete dataset for this study will be placed in the Carefish/catch project webpage https://carefish.net/catch upon publication.

## Suplemental data

**Table S1:**
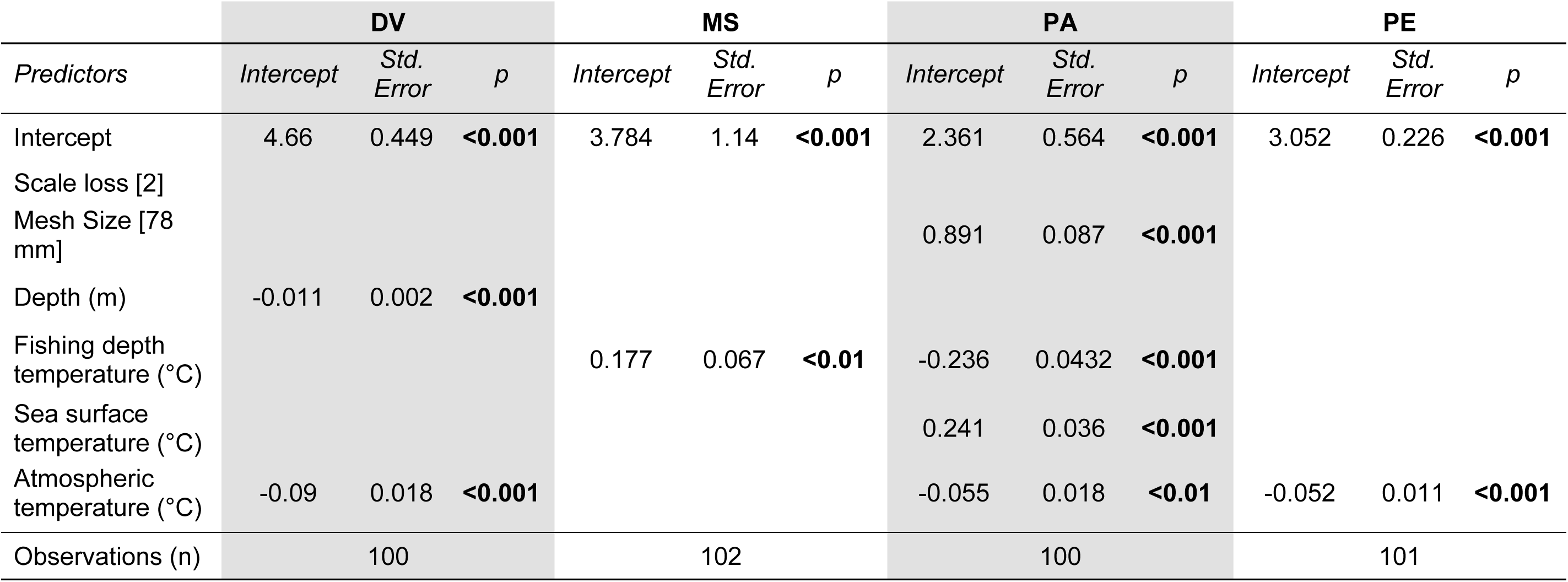
The Estimate, standard error, and the p-values (p: Pr(>|z|)) derived from the Generalised linear models (GLMs) that were fit to predict the impact of several biological, operational, and environmental predictors on the duration of fish activity on the deck of the fishing vessel. (DV: Two-banded seabream (*Diplodus vulgaris*), MS: Red mullet (*Mullus surmuletus*), PA: Axillary seabream (*Pagellus acarne*), PE: Common pandora (*Pagellus erythrinus*).

|  | DV |  |  | MS |  |  | PA |  |  | PE |  |  |
| --- | --- | --- | --- | --- | --- | --- | --- | --- | --- | --- | --- | --- |
| Predictors | Intercept | Std. Error | p | Intercept | Std. Error | p | Intercept | Std. Error | p | Intercept | Std. Error | p |
| Intercept | 4.66 | 0.449 | <0.001 | 3.784 | 1.14 | <0.001 | 2.361 | 0.564 | <0.001 | 3.052 | 0.226 | <0.001 |
| Scale loss [2] |  |  |  |  |  |  |  |  |  |  |  |  |
| Mesh Size [78 mm] |  |  |  |  |  |  | 0.891 | 0.087 | <0.001 |  |  |  |
| Depth (m) | -0.011 | 0.002 | <0.001 |  |  |  |  |  |  |  |  |  |
| Fishing depth temperature (°C) |  |  |  | 0.177 | 0.067 | <0.01 | -0.236 | 0.0432 | <0.001 |  |  |  |
| Sea surface temperature (°C) |  |  |  |  |  |  | 0.241 | 0.036 | <0.001 |  |  |  |
| Atmospheric temperature (°C) | -0.09 | 0.018 | <0.001 |  |  |  | -0.055 | 0.018 | <0.01 | -0.052 | 0.011 | <0.001 |
| Observations (n) | 100 |  |  | 102 |  |  | 100 |  |  | 101 |  |  |

**Table S2:** The Odds ratios, CI: Confidence intervals, and the p-values derived from the Cumulative linked mixed models (CLMMs) that were fit to predict the impact of several biological, operational, and environmental predictors on the vitality at the time of landing on deck. (All species: The full model consisting of all the species, DV: Two-banded seabream (*Diplodus vulgaris*), MS: Red mullet (*Mullus surmuletus*), PA: Axillary seabream (*Pagellus acarne*); Vitality scale: 4 = highly active, 3 = less active, 2 = lethargic and 1 = unresponsive).

|  | All species |  |  | DV |  |  | MS |  |  | PA |  |  |
| --- | --- | --- | --- | --- | --- | --- | --- | --- | --- | --- | --- | --- |
| Predictors | Odds Ratios | CI | p | Odds Ratios | CI | p | Odds Ratios | CI | p | Odds Ratios | CI | p |
| 1 \| 2 | 0.05 | 0.03-0.11 | **<0.001** | 0.01 | 0.00-0.04 | **<0.001** | 1.08 | 0.43-2.75 | 0.865 | 0.12 | 0.05-0.27 | **<0.001** |
| 2 \| 3 | 0.21 | 0.11-0.39 | **<0.001** | 0.08 | 0.03-0.25 | **<0.001** | 4.32 | 1.60-11.67 | **0.004** | 0.67 | 0.37-1.21 | 0.182 |
| 3 \| 4 | 0.77 | 0.42-1.42 | 0.4 | 1.05 | 0.44-2.53 | 0.905 | 8.79 | 3.10-24.95 | **<0.001** | 1.49 | 0.85-2.61 | 0.16 |
| Species [MS] | 0.08 | 0.04-0.17 | **<0.001** |  |  |  |  |  |  |  |  |  |
| Species [PA] | 1.01 | 0.51-2.01 | 0.97 |  |  |  |  |  |  |  |  |  |
| Species [PE] | 1.68 | 0.85-3.32 | 0.138 |  |  |  |  |  |  |  |  |  |
| Scale loss [2] | 0.27 | 0.11-0.65 | **0.003** | 0.02 |  | **0.039** | 0.36 |  | 0.077 |  |  |  |
| Length (cm) |  |  |  |  |  |  | 0.52 |  | **0.028** |  |  |  |
| Mesh size [78mm] |  |  |  |  |  |  | 7.54 |  | 0.137 | 20.02 | 4.71-85.06 | **<0.001** |
| Fishing depth temperature (°C) |  |  |  |  |  |  | 0.4 |  | **0.033** |  |  |  |
| Sea surface temperature (°C) |  |  |  | 2.2 | 1.03-4.69 | **0.042** | 0.0 |  |  | 2.6 | 1.53-4.41 | **<0.001** |
| Observations (n) | 403 |  |  | 100 |  |  | 100 |  |  | 102 |  |  |

**Table S3:**
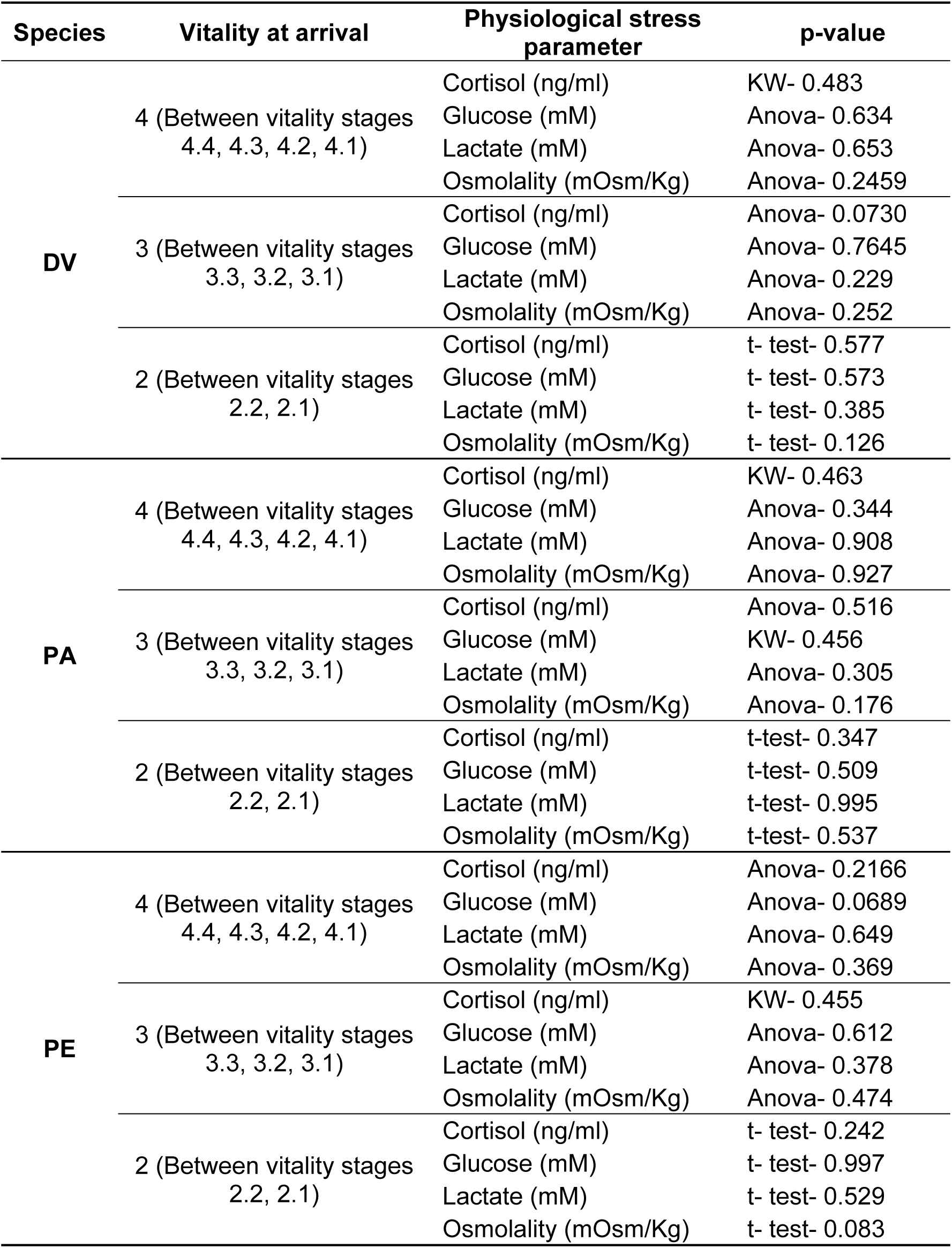
The p-values obtained from statistical significance tests performed to compare the levels of physiological stress parameters between vitality stages depending upon the vitality at arrival (DV: Two-banded seabream (*Diplodus vulgaris*), PA: Axillary seabream (*Pagellus acarne*), PE: *Pagellus erythrinus*; Anova: Analysis of variance, KW: Kruskal-Wallis test, t-test: Welch two sample t-test).

| Species | Vitality at arrival | Physiological stress parameter | p-value |
| --- | --- | --- | --- |
| DV | 4 (Between vitality stages 4.4, 4.3, 4.2, 4.1) | Cortisol (ng/ml) | KW- 0.483 |
|  |  | Glucose (mM) | Anova- 0.634 |
|  |  | Lactate (mM) | Anova- 0.653 |
|  |  | Osmolality (mOsm/Kg) | Anova- 0.2459 |
|  | 3 (Between vitality stages 3.3, 3.2, 3.1) | Cortisol (ng/ml) | Anova- 0.0730 |
|  |  | Glucose (mM) | Anova- 0.7645 |
|  |  | Lactate (mM) | Anova- 0.229 |
|  |  | Osmolality (mOsm/Kg) | Anova- 0.252 |
|  | 2 (Between vitality stages 2.2, 2.1) | Cortisol (ng/ml) | t- test- 0.577 |
|  |  | Glucose (mM) | t- test- 0.573 |
|  |  | Lactate (mM) | t- test- 0.385 |
|  |  | Osmolality (mOsm/Kg) | t- test- 0.126 |
| PA | 4 (Between vitality stages 4.4, 4.3, 4.2, 4.1) | Cortisol (ng/ml) | KW- 0.463 |
|  |  | Glucose (mM) | Anova- 0.344 |
|  |  | Lactate (mM) | Anova- 0.908 |
|  |  | Osmolality (mOsm/Kg) | Anova- 0.927 |
|  | 3 (Between vitality stages 3.3, 3.2, 3.1) | Cortisol (ng/ml) | Anova- 0.516 |
|  |  | Glucose (mM) | KW- 0.456 |
|  |  | Lactate (mM) | Anova- 0.305 |
|  |  | Osmolality (mOsm/Kg) | Anova- 0.176 |
|  | 2 (Between vitality stages 2.2, 2.1) | Cortisol (ng/ml) | t-test- 0.347 |
|  |  | Glucose (mM) | t-test- 0.509 |
|  |  | Lactate (mM) | t-test- 0.995 |
|  |  | Osmolality (mOsm/Kg) | t-test- 0.537 |
| PE | 4 (Between vitality stages 4.4, 4.3, 4.2, 4.1) | Cortisol (ng/ml) | Anova- 0.2166 |
|  |  | Glucose (mM) | Anova- 0.0689 |
|  |  | Lactate (mM) | Anova- 0.649 |
|  |  | Osmolality (mOsm/Kg) | Anova- 0.369 |
|  | 3 (Between vitality stages 3.3, 3.2, 3.1) | Cortisol (ng/ml) | KW- 0.455 |
|  |  | Glucose (mM) | Anova- 0.612 |
|  |  | Lactate (mM) | Anova- 0.378 |
|  |  | Osmolality (mOsm/Kg) | Anova- 0.474 |
|  | 2 (Between vitality stages 2.2, 2.1) | Cortisol (ng/ml) | t- test- 0.242 |
|  |  | Glucose (mM) | t- test- 0.997 |
|  |  | Lactate (mM) | t- test- 0.529 |
|  |  | Osmolality (mOsm/Kg) | t- test- 0.083 |

**Table S4:**
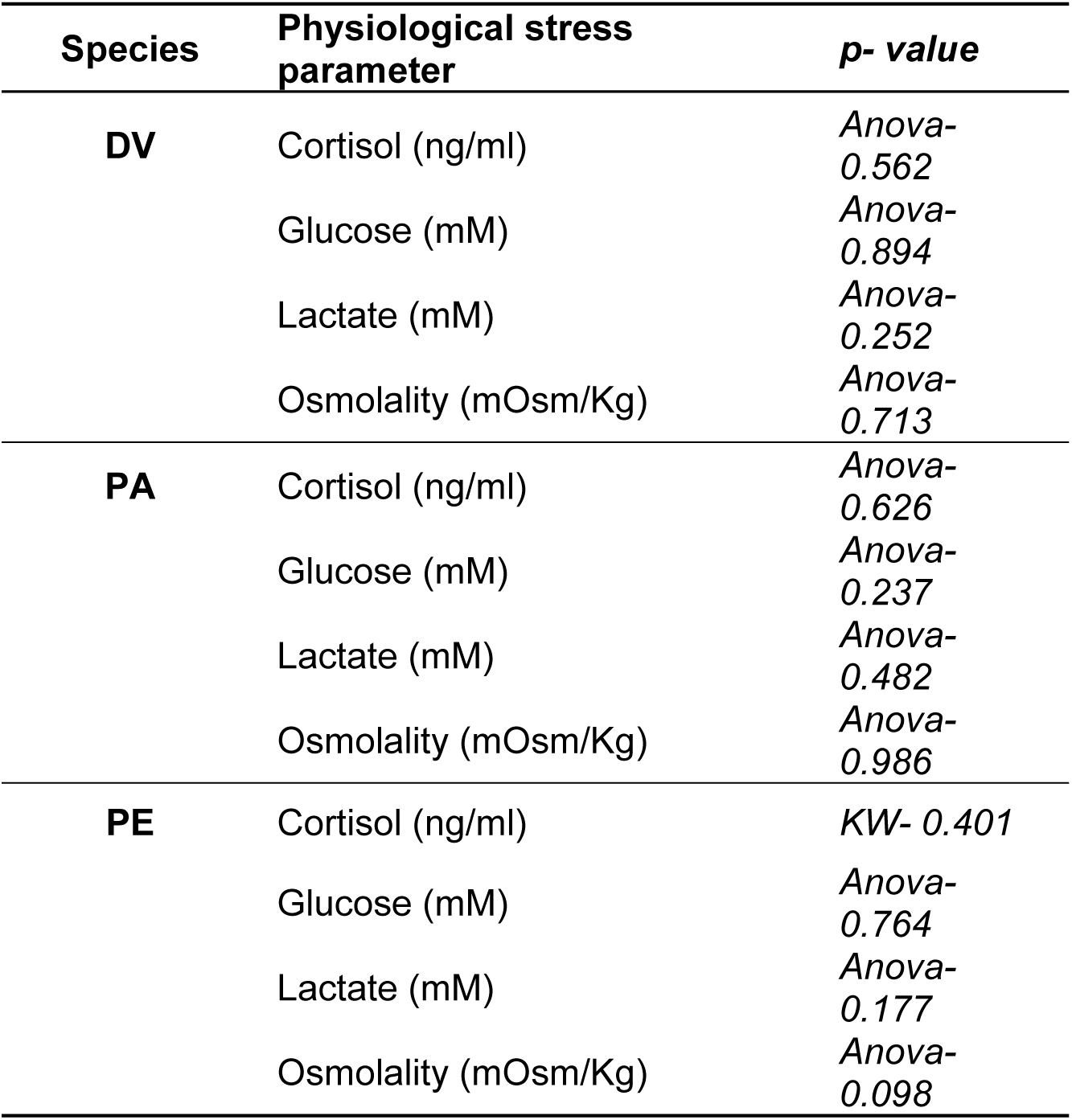
The p-values obtained from statistical significance tests performed to compare the levels of physiological stress parameters in vitality stages where the vitality at arrival was the same as vitality at sampling (stages 4.4, 3.3, 2.2, and 1.1) as indicated by the striped bars in Figure 7. (DV: Two-banded seabream (*Diplodus vulgaris*), PA: Axillary seabream (*Pagellus acarne*), PE: *Pagellus erythrinus*; Anova: Analysis of variance, KW: Kruskal-Wallis test).

| Species | Physiological stress parameter | p- value |
| --- | --- | --- |
| DV | Cortisol (ng/ml) | Anova- 0.562 |
|  | Glucose (mM) | Anova- 0.894 |
|  | Lactate (mM) | Anova- 0.252 |
|  | Osmolality (mOsm/Kg) | Anova- 0.713 |
| PA | Cortisol (ng/ml) | Anova- 0.626 |
|  | Glucose (mM) | Anova- 0.237 |
|  | Lactate (mM) | Anova- 0.482 |
|  | Osmolality (mOsm/Kg) | Anova- 0.986 |
| PE | Cortisol (ng/ml) | KW- 0.401 |
|  | Glucose (mM) | Anova- 0.764 |
|  | Lactate (mM) | Anova- 0.177 |
|  | Osmolality (mOsm/Kg) | Anova- 0.098 |

